# Poly(2-oxazoline)-based polyplexes as a PEG-free plasmid DNA delivery platform

**DOI:** 10.1101/2022.12.18.518592

**Authors:** Dina N. Yamaleyeva, Naoki Makita, Duhyeong Hwang, Matthew J. Haney, Rainer Jordan, Alexander V. Kabanov

## Abstract

The present study expands the versatility of cationic poly(2-oxazoline) (POx) copolymers as a PEG-free platform for gene delivery to immune cells, such as monocytes and macrophages. Several block copolymers are developed by varying non-ionic hydrophilic blocks (poly(2-methyl-2-oxazoline) (pMeOx) or poly(2-ethyl-2-oxazoline) (pEtOx), cationic blocks, and an optional hydrophobic block (poly(2-isopropyl-2-oxazoline) (iPrOx). The cationic blocks are produced by side chain modification of 2-methoxy-carboxyethyl-2-oxazoline (MestOx) block precursor with diethylenetriamine (DET) or tris(2-aminoethyl)amine (TREN). For the attachment of a targeting ligand, mannose, we employed azide-alkyne cycloaddition click chemistry methods. Of the two cationic side chains, polyplexes made with DET-containing copolymers transfect macrophages significantly better than those made with TREN-based copolymer. Likewise, non-targeted pEtOx-based diblock copolymer is more active in cell transfection than pMeOx-based copolymer. The triblock copolymer with hydrophobic block iPrOx performs poorly compared to the diblock copolymer which lacks this additional block. Surprisingly, attachment of a mannose ligand to either of these copolymers is inhibitory for transfection. Despite similarities in size and design, mannosylated polyplexes result in lower cell internalization compared to non-mannosylated polyplexes. Thus, PEG-free, non-targeted DET- and pEtOx-based diblock copolymer outperforms other studied structures in the transfection of macrophages and displays transfection levels comparable to GeneJuice, a commercial non-lipid transfection reagent.

## 1. Introduction

Despite dramatic progress in application of the lipid nanoparticles (LNP) in mRNA vaccines the gene delivery systems face challenging barriers such as off-target immune responses and low delivery efficiency. Genetic material, such as plasmid DNA (pDNA), is challenging to deliver *in vivo* due to degradation by DNases, inefficient delivery into the cell, and lysosomal entrapment and degradation.^[1]^ Polymer-mediated gene delivery focuses on combining cationic polymers with negatively charged genetic material to form polyion complexes (polyplexes). One of the very well-studied cationic polymers used for plasmid delivery is a block copolymer of polyethyleneimine (PEI) and polyethylene glycol (PEG).^[1,2]^ Though PEI-PEG based polyplexes have a high transfection efficiency, the high molecular weight of PEI is required for efficient transfection, which makes these polyplexes cytotoxic and unsuitable for *in vivo* application.^[2,3]^ The addition of PEG results in a beneficial “stealth” effect allowing for enhanced circulation *in vivo*.^[4,5]^ Due to its relative inertness and “stealth” property, PEG quickly became used in many cancer treatments, such as in breast and ovarian cancer drugs. However, the ubiquity of PEG is problematic in causing the rise of PEG-antibodies, which decreases the efficacy of life-saving PEG-based treatments.^[5]^ A recent study reports that 72% of individuals have detectable levels of PEG antibodies, which is driving a significant need for alternative polymers that also employ stealth properties.^[6]^ One promising candidate for PEG replacement is poly(2-oxazoline), or POx.

Poly(2-oxazolines) are a new and alternative class of polymer compared to PEG with numerous advantages.^[7,8]^ Our group has proven that POx has many convenient features such as adjustable hydrophobicity and straightforward chemistry allowing for precise customization of block orientation and block lengths.^[9–11]^ Two hydrophilic POx monomers, 2-methyl-2-oxazoline (MeOx) and 2-ethyl-2-oxazoline (EtOx), stand out as suitable alternatives to PEG due to their similar biocompatibility, stability *in vivo*, and reduced production of reactive oxygen species.^[12]^. Our team has successfully used amphiphilic POx block copolymers to improve capacity of synergistic drug combinations as well as solubilized previously insoluble drugs with high efficiency.^[13,14]^ We have also reported on the POx cationic block copolymer used to formulate pDNA into polyplexes with decreased serum binding compared to PEG-based polyplexes. Due to this success in past works, the present study focuses on further expanding the versatility of POx block copolymers by developing a platform for pDNA delivery to immune cells, such as monocytes and macrophages. Specifically, our polymers are modified with a moiety targeting the macrophage mannose receptor (MMR). Macrophages are a natural target as they have been implicated in worsening cancer progression.^[15–17]^ The mannose receptor was chosen as a targeting ligand because of its ubiquity on the surface of macrophages and reported ability to enhance uptake and therefore transfection.^[18]^ Studies estimate that 20-70% of a breast cancer tumor can be composed of macrophages and tumor-associated macrophages.^[19]^ By successfully transfecting macrophages with a PEG-free gene delivery system, future POx-based breast cancer treatments can be developed.

To design an optimized polymer for macrophage transfection, various configurations of POx-based block copolymers for pDNA delivery were designed and compared for their transfection efficacy. By taking advantage of the various properties of each block, a series of diblock and triblock polymers were investigated by analyzing the following: 1) the comparison between MeOx and EtOx monomers, 2) the effect of two cationic side chain modifications using diethylenetriamine (DET) (linear) or tris(2-aminoethyl)amine (TREN) (branched), 3) effect of conjugating the targeting moiety, mannose, via copper-catalyzed azide-alkyne cycloaddition (CuAAC) click chemistry, and 4) how an additional hydrophobic piPrOx block affects the structure, stability, and transfection ability of luciferase-encoding pDNA (luc-pDNA) polyplexes. By considering each of the properties of these modifications, we aim to develop a non-toxic PEG-free transfection platform, which has a high transfection efficiency in various immune cell lines. In this study, we focus on POx-pDNA polyplexes as a PEG-free alternative for pDNA delivery to immune cells such as macrophages.

## 2. Results

### 2.1. Synthesis of cationic poly(2-oxazoline) block copolymers

To design a PEG-free polymer for plasmid transfection of immune cells, such as macrophages, we developed several POx-based cationic copolymers by varying non-ionic hydrophilic, cationic, and hydrophobic blocks, and employing azide-alkyne cycloaddition (“click chemistry”) methods for the attachment of the targeting moiety **(Figure 1)**. The polymers were synthesized by sequential living cationic ring-opening polymerization (LCROP) of 2-oxazoline monomers which provides access to a wide range of polymer structures with defined molecular mass and narrow dispersity (1.01-1.30).^[20]^ We used two different strategies to introduce “clickable” groups for targeting moieties attachment to the free ends of the hydrophilic blocks. In one strategy **(Figure 2A)** we employed p-toluenesulfonic acid methyl ester as the initiator, and first polymerized the cationic block precursor pMestOx, followed by the hydrophilic block, which was terminated by DBCO-amine for copper-free click chemistry. In the second strategy **(Figure 2B)**, we employed alkyne-containing propargyl p-toluenesulfonate as the initiator and then sequentially polymerized the hydrophilic block and the cationic block precursor that was terminated by piperidine. We varied the hydrophilic block structure using either MeOx or EtOx as the monomers (Figure 2A). We also introduced a third type of relatively hydrophobic block by polymerizing iPrOx after the cationic precursor (Figure 2B), to enhance block copolymer self-assembly during polyplex formation. After synthesis of the block copolymer precursor, we attached the mannose targeting moiety Alpha-Mann-TEG-N_3_ using copper-free (Figure 2A) or copper-catalyzed (Figure 2B) click chemistry. Finally, we introduced the cationic moieties by reacting the methyl ester groups of the corresponding block copolymer precursors with either DET or TREN. The resulting polymers are presented in **Table 1**. Mannose conjugation was confirmed via NMR **(Supplementary Figure S1, S2)**. Polymers were characterized with 2-(p-toluidino)-6-naphthalene sulfonic acid (TNS) assay, pH titration, and by examining buffering capacity **(Supplementary Figure S3 C – E)**. We further used the TNS assay to examine the DET and TREN-containing MeOx block copolymers. TNS fluorescence increases upon binding to the protonated amines. The fluorescence intensity of TNS upon mixing with the TREN-containing copolymer was constant across the pH 4.0 to pH 7.4 range, which suggests that the TREN side chains were protonated in this range pH. In contrast for the DET-containing copolymer, the fluorescence signal increased at lower upon acidification from pH 5.0 to pH 4.0, with is indicative of the amino group protonation in this range (Supplementary Figure S3 C). The pH titration study suggests that these polymers display buffering capacity in a broad range of pH indicative of protonation of multiple amino groups. Specifically, the methyl-based DET-containing copolymer displays buffering capacity in both acidic and alkali areas with effective pKa values of approximately 6.0 and 11.0, while the TREN-containing polymer displayed a buffering capacity in the ranges corresponding to effective pKa of approximately 4.0 and 10.0. (Supplementary Figure S3D and S3E). Ethyl-based DET-containing copolymer also displayed a buffering capacity in both acidic and alkali regions with effective pKa close to 4.3 to 8.8 respectively **(Supplementary Figure S4)**.

**Table 1.**
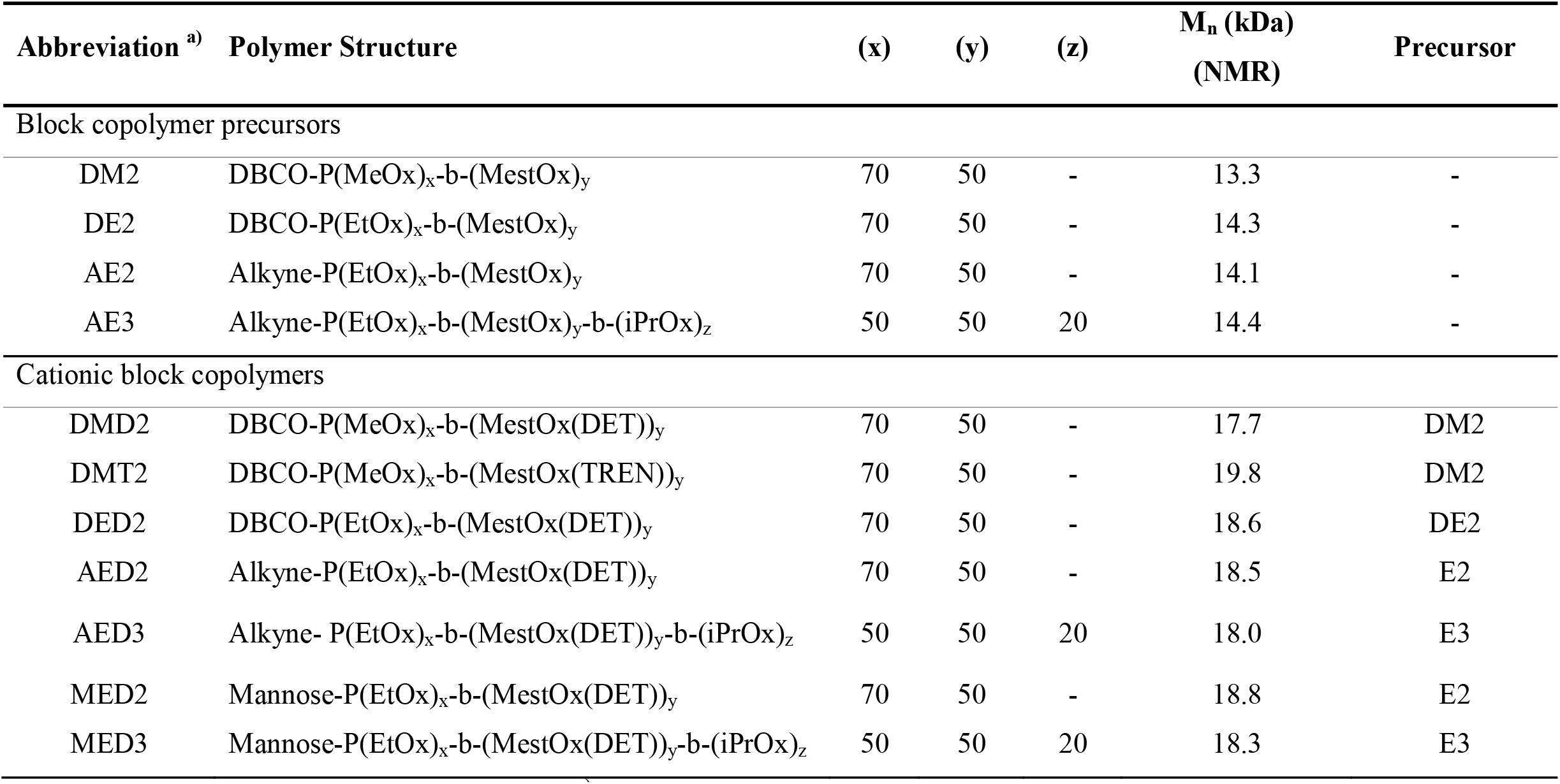
Polymers are code named as follows.^a)^ In the two letter codes, the first letter represents DBCO (D) or Alkyne (A). The second letter represents hydrophilic non-ionic block, MeOx (M) or EtOx (E). In the three letter codes, the first letter represents DBCO (D), Alkyne (A), or mannose (M). The second letter represents hydrophilic non-ionic block, MeOx (M) or EtOx (E), and the third letter represents the cationic block DET (D) or TREN (T). In all cases and the concluding number stands for diblock (2) or triblock (3) copolymer structure.

| Abbreviation <sup>a)</sup> | Polymer Structure | (x) | (y) | (z) | M<sub>n</sub> (kDa)<br>(NMR) | Precursor |
| --- | --- | --- | --- | --- | --- | --- |
| Block copolymer precursors |  |  |  |  |  |  |
| DM2 | DBCO-P(MeOx)<sub>x</sub>-b-(MestOx)<sub>y</sub> | 70 | 50 | - | 13.3 | - |
| DE2 | DBCO-P(EtOx)<sub>x</sub>-b-(MestOx)<sub>y</sub> | 70 | 50 | - | 14.3 | - |
| AE2 | Alkyne-P(EtOx)<sub>x</sub>-b-(MestOx)<sub>y</sub> | 70 | 50 | - | 14.1 | - |
| AE3 | Alkyne-P(EtOx)<sub>x</sub>-b-(MestOx)<sub>y</sub>-b-(iPrOx)<sub>z</sub> | 50 | 50 | 20 | 14.4 | - |
| Cationic block copolymers |  |  |  |  |  |  |
| DMD2 | DBCO-P(MeOx)<sub>x</sub>-b-(MestOx(DET))<sub>y</sub> | 70 | 50 | - | 17.7 | DM2 |
| DMT2 | DBCO-P(MeOx)<sub>x</sub>-b-(MestOx(TREN))<sub>y</sub> | 70 | 50 | - | 19.8 | DM2 |
| DED2 | DBCO-P(EtOx)<sub>x</sub>-b-(MestOx(DET))<sub>y</sub> | 70 | 50 | - | 18.6 | DE2 |
| AED2 | Alkyne-P(EtOx)<sub>x</sub>-b-(MestOx(DET))<sub>y</sub> | 70 | 50 | - | 18.5 | E2 |
| AED3 | Alkyne- P(EtOx)<sub>x</sub>-b-(MestOx(DET))<sub>y</sub>-b-(iPrOx)<sub>z</sub> | 50 | 50 | 20 | 18.0 | E3 |
| MED2 | Mannose-P(EtOx)<sub>x</sub>-b-(MestOx(DET))<sub>y</sub> | 70 | 50 | - | 18.8 | E2 |
| MED3 | Mannose-P(EtOx)<sub>x</sub>-b-(MestOx(DET))<sub>y</sub>-b-(iPrOx)<sub>z</sub> | 50 | 50 | 20 | 18.3 | E3 |

**Figure 1.**
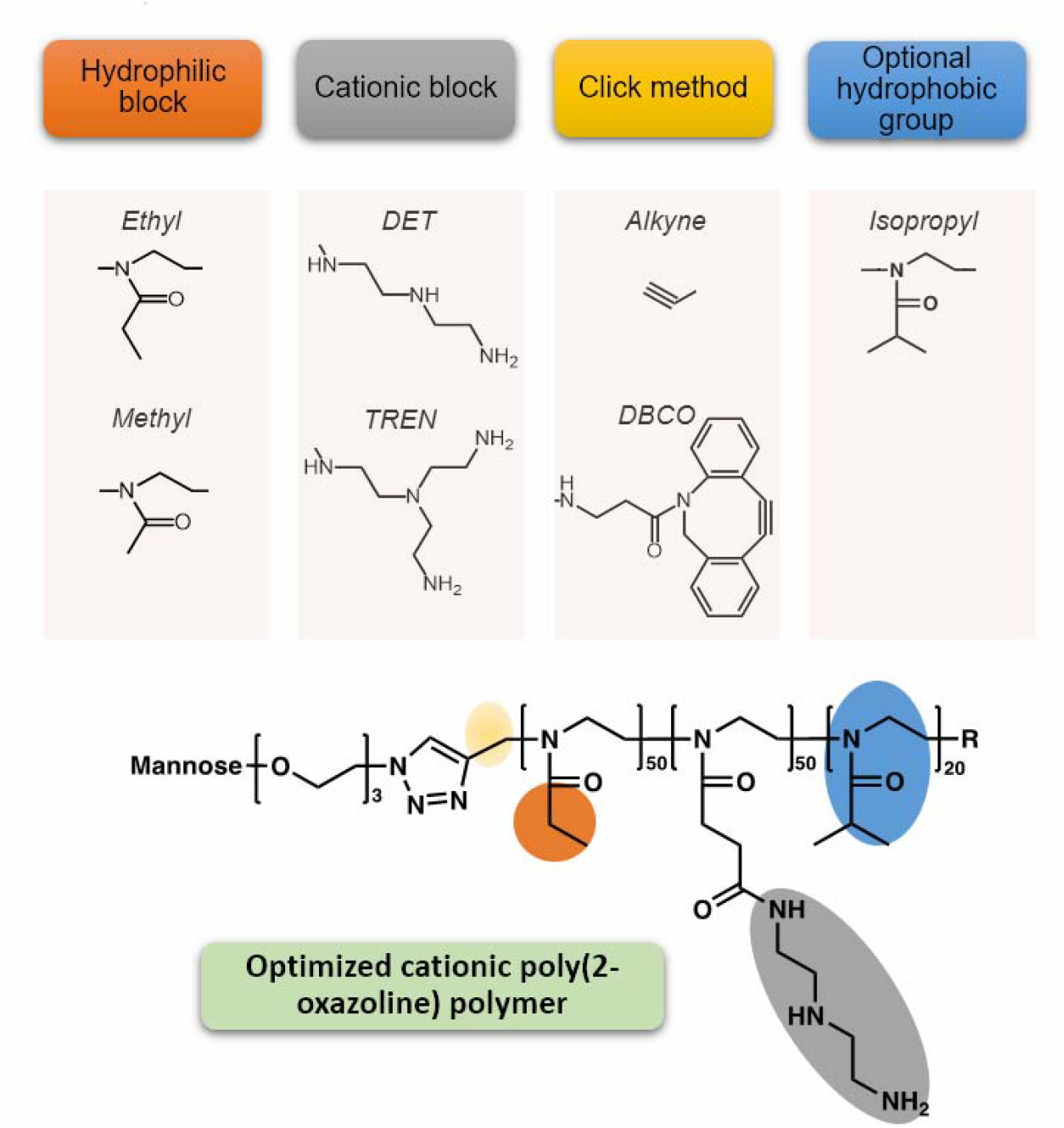
Scheme. Overall scheme of optimization strategy for a targeted poly(2-oxazoline)-based polymer capable of transfecting immune cells, such as macrophages, via polyplex formation with plasmid DNA. Components varied in polymer design include the following groups: hydrophilic block, cationic block, copper-free or copper-based click method for conjugating mannose targeting moiety, and an optional thermosensitive-hydrophobic group.

**Figure 2.**
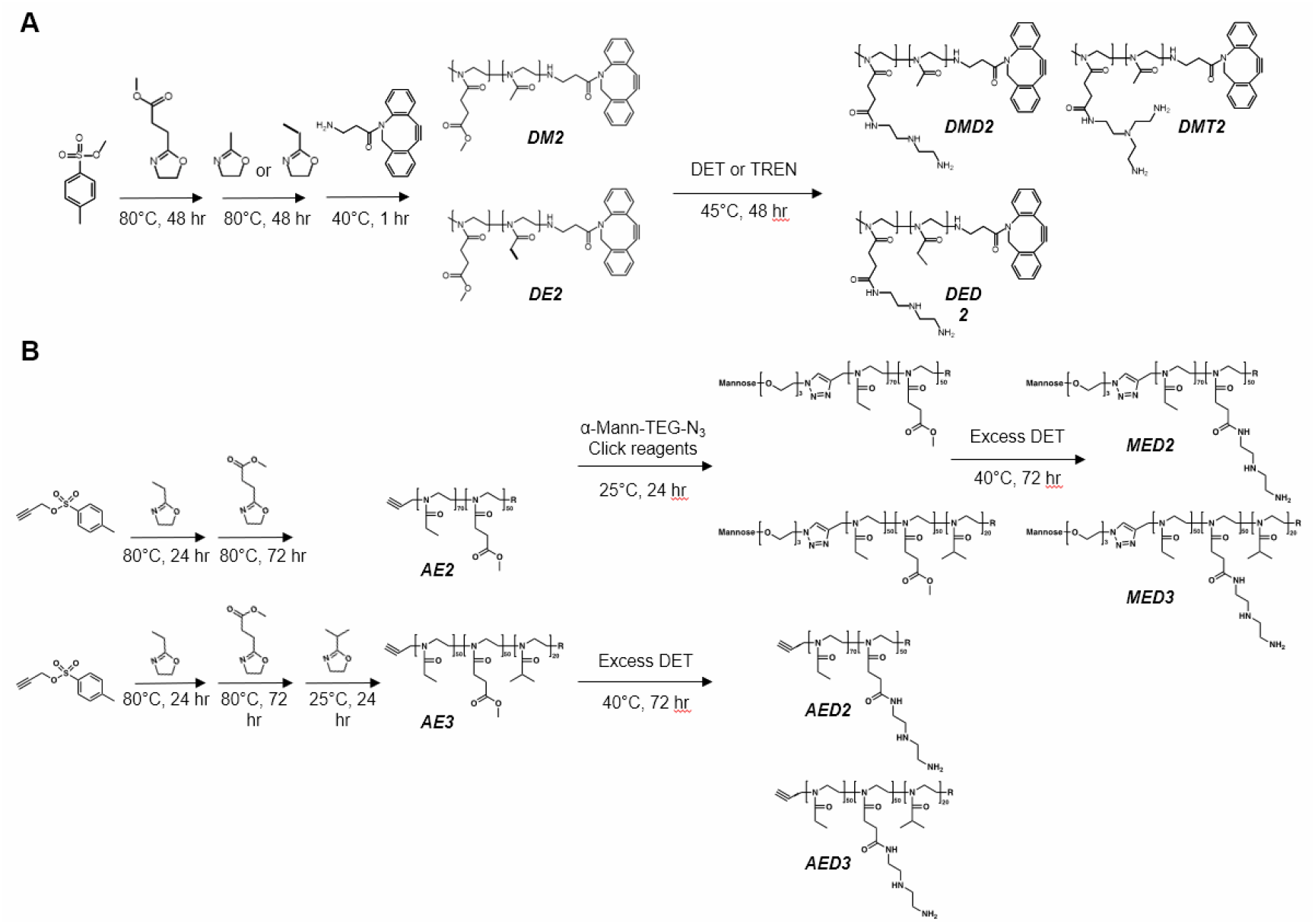
Synthesis schemes. We designed two synthetic strategies for development of cationic polymers as gene delivery vehicles with a clickable moiety for further modification. (A) A diblock copolymer composed of pMestOx and pEtOx was synthesized via sequential LCROP of 2-oxazolines initiated by p-toluenesulfonic acid methyl ester and terminated with DBCO-amine. Diethylenetriamine (DET) or tris(2-aminoethyl)amine (TREN) was incorporated to the methylester group of the diblock copolymer via ester-amide exchange reaction. Resulting polymers were DMD2, DMT2, or DED2. (B) Diblock and triblock copolymers composed of pEtOx, pMestOx, and piPrOx (triblock only) were synthesized via sequential LCROP with propargyl p-toluenesulfonate as the initiator and terminated with piperidine (R). DET was incorporated into the methylester group via ester-amide exchange reaction. Polymers were also mannosylated resulting in four final polymers consisting of EtOx, DET, and optional mannose: AED2, MED2, AED3, and MED3.

### 2.2. Formation of polyplexes depends on polycation structure and mannosylation

To produce the polyion complexes the cationic copolymers were mixed with luciferase-encoding pDNA using simple vortex mixing at various N/P ratios and incubated at room temperature for 30 minutes prior to any characterization. To obtain different N/P ratios the amount of luc-pDNA was kept constant (33 mg/mL) while the concentration of the polymer was varied in each polyplex formulation. The formation of the polyplexes was detected by the changes of the electrophoretic mobility of the luc-pDNA in 1% agarose gel, by particle size measurements using dynamic light scattering (DLS) as well as TEM **(Figure 3 A-C and Supplementary Figure S3 A)**. Generally, the particle sizes for polyplexes of various compositions varied from ca. 70 to ca. 120 nm with fairly narrow polydispersity index (PDI ca. 0.2) **(Supplementary Figure S3 B and Supplementary Figure S5)**. To examine the morphology, polyplexes were prepared at N/P 20 and then imaged with TEM. The complexes were distinct, non-aggregated and either spherical or somewhat elongated (short worms) **(Figure 3C)**. No difference in size was observed between polymers with MeOx block compared to EtOx block (Supplementary Figure S3B). Thus, at lower N/P ratios the DET containing diblock copolymers displayed some disproportioning - i.e., presence of free luc-pDNA or negatively charged complexes that were mobile in gels, along with the polyplexes remaining at the start of the gel **(Supplementary Figure S6)**. The TREN containing diblock copolymers revealed greater propensity for formation of the complexes than the DET containing copolymers (Supplementary Figure S6). This probably was due to higher charge density of the TREN (three chargeable amino groups) vs DET (two chargeable amino groups). Addition of the third hydrophobic piPrOx block in the copolymer increased the tendency for disproportioning. The triblock copolymer **AED3** (P(EtOx)_50_-b-(MestOx(DET))_50_-b-(iPrOx)_20_-Alkyne) did not form complexes well at lower N/P ratios of 1 and 2, although the complexation at N/P 10 and 20 was nearly complete (**Figure 3B** and Supplementary Figure S6). Another factor impacting copolymer binding to the luc-pDNA was attachment of the mannose. At lower N/P ratios of 1 and 2 this appeared to hinder the polyion complexation of mannosylated block copolymers with pDNA in contrast to the non-mannosylated counterparts (Supplementary Figure S6). The least effective binding was observed with the mannosylated triblock **MED3** (P(EtOx)_50_-b-(MestOx(DET))_50_-b- (iPrOx)_20_-Mannose). For this copolymer, some mobility of the luc-pDNA in the gel was seen even at highest N/P ratios 10 and 20 (Figure 3B). With all copolymers at such high N/P ratios the polyplexes were positively charged as follows from the zeta-potential measurements **(Supplementary Figure S7)**. To understand the complexation of mannosylated polymers further, polyplexes were tested by an ethidium bromide (EtBr) displacement assay. By forming polyplexes with a mixture of EtBr and luc-pDNA, the polymer competes against EtBr which allows us to monitor luc-pDNA condensation. Though gel electrophoresis showed a lack of complexation at lower N/P ratios of 1 and 2 for polyplexes based on both diblock and triblock copolymers, EtBr displacement revealed that these same polyplexes displaced EtBr successfully starting at N/P ratio 2 **(Figure 5A)**. Both diblock-based and triblock-based polyplexes showed similar EtBr displacement levels at ∼85% displacement (Figure 5A). Interestingly, polymer mannosylation did not affect the amount of EtBr displaced.

**Figure 3.**
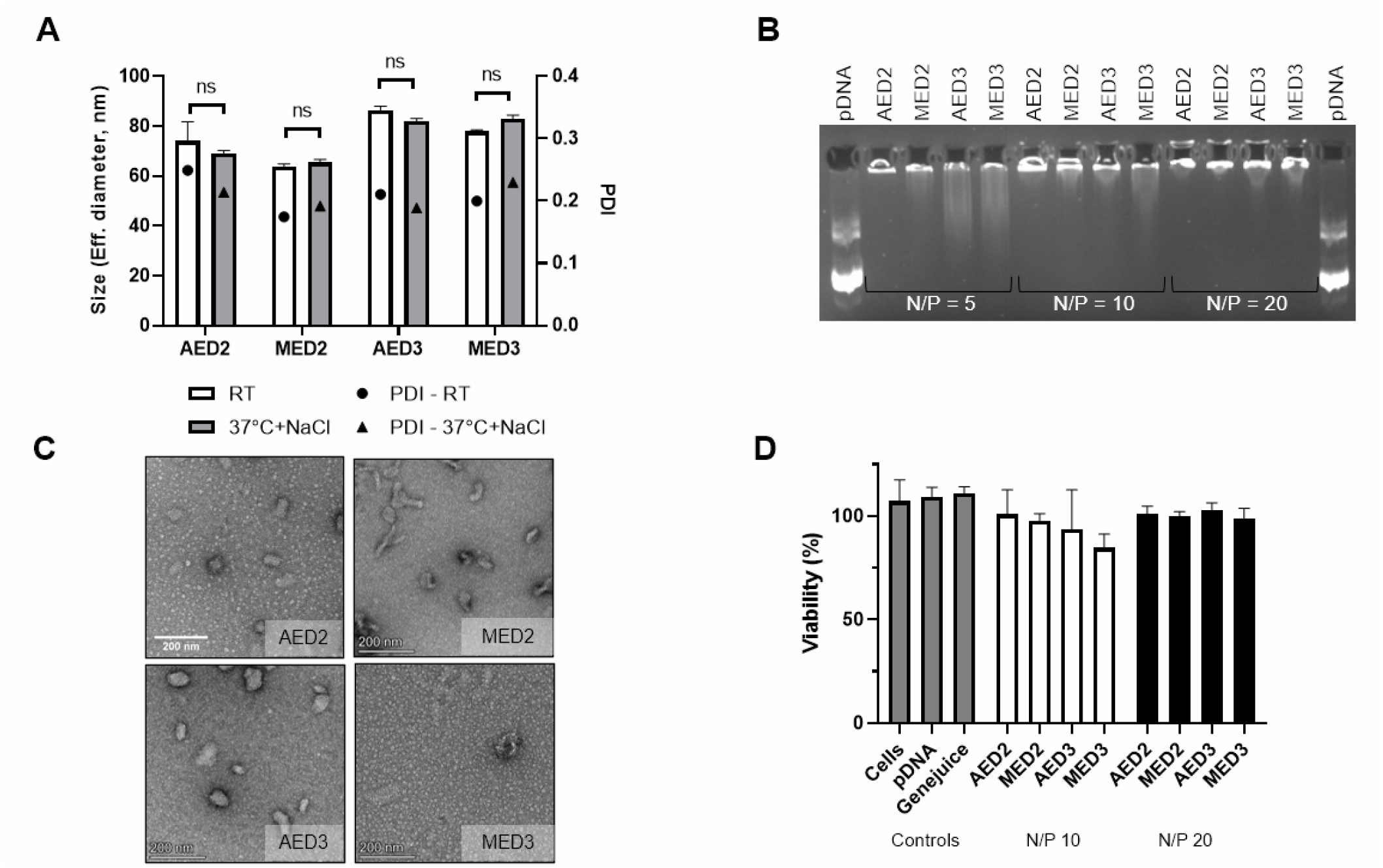
Polyplex characterization. (A) Stability of polyplexes by size (bars) and polydispersity index (PDI) (symbols) at N/P 20 after 30 min incubation at room temperature (RT) or after 30 min incubation at room temperature followed by a 60 min incubation at 37°C in 150 mM NaCl (37°C+NaCl). (B) Gel electrophoresis showing complexation between polymers and luc-pDNA at NP ratios 5, 10, and 20 after 30 min incubation at RT. (C) TEM images of AED2, MED2, AED3, or MED3-based polyplexes prepared at N/P 20 after 30 min incubation at RT. (D) RAW264.7 viability after 24 hour treatment with polyplexes prepared at N/P 10 and 20. ns: not significant.

**Figure 4.**
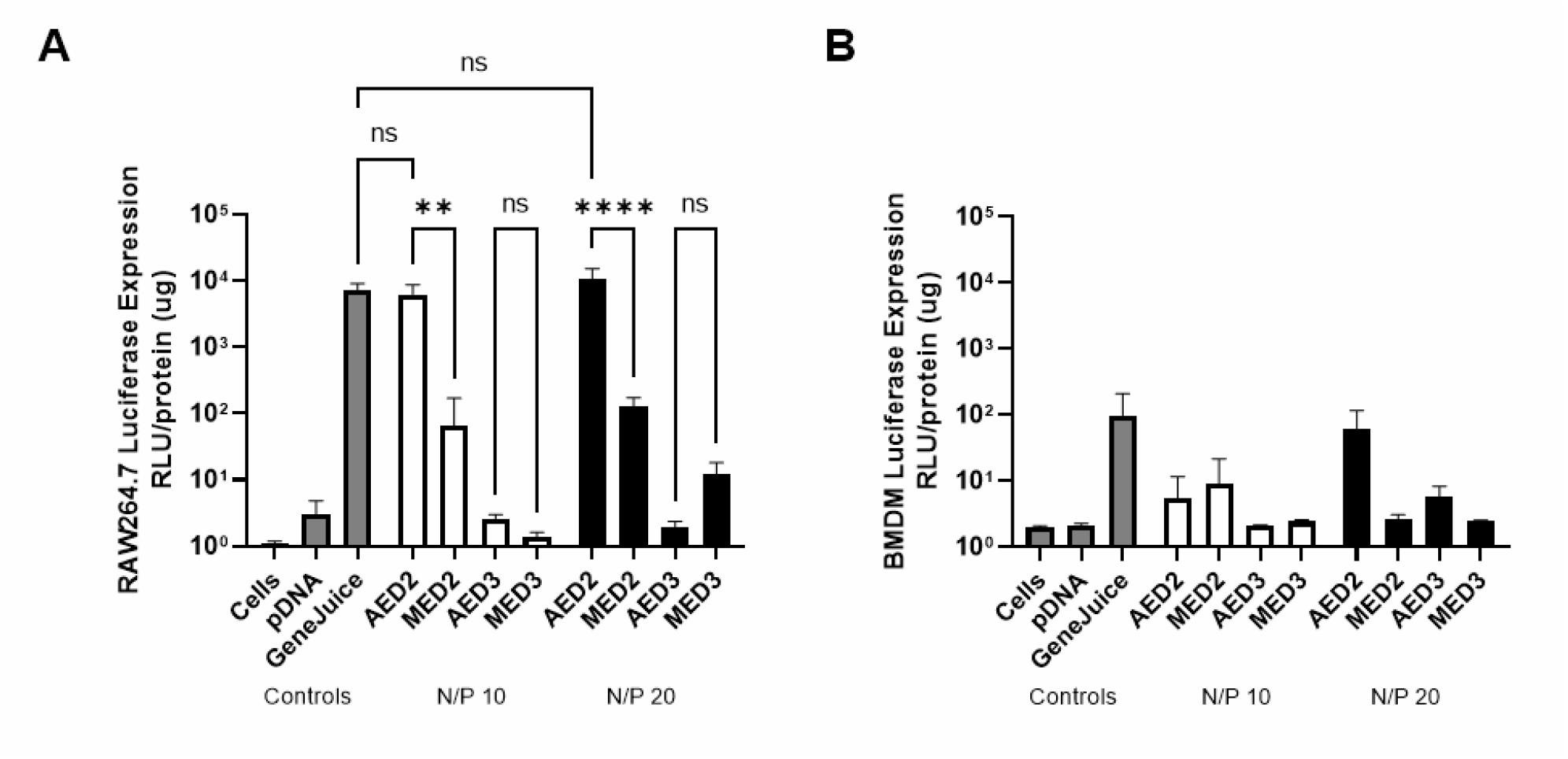
*In vitro* transfection screening. Screening macrophage transfection with optimized targeted and untargeted diblock and triblock polymers in (A) RAW264.7 and (B) BMDM. Controls are cells alone, pDNA alone, and GeneJuice, a commercial transfection reagent. ns: not significant. *p<0.5, **p<0.01, ***p<0.001, ****p<0.0001.

**Figure 5.**
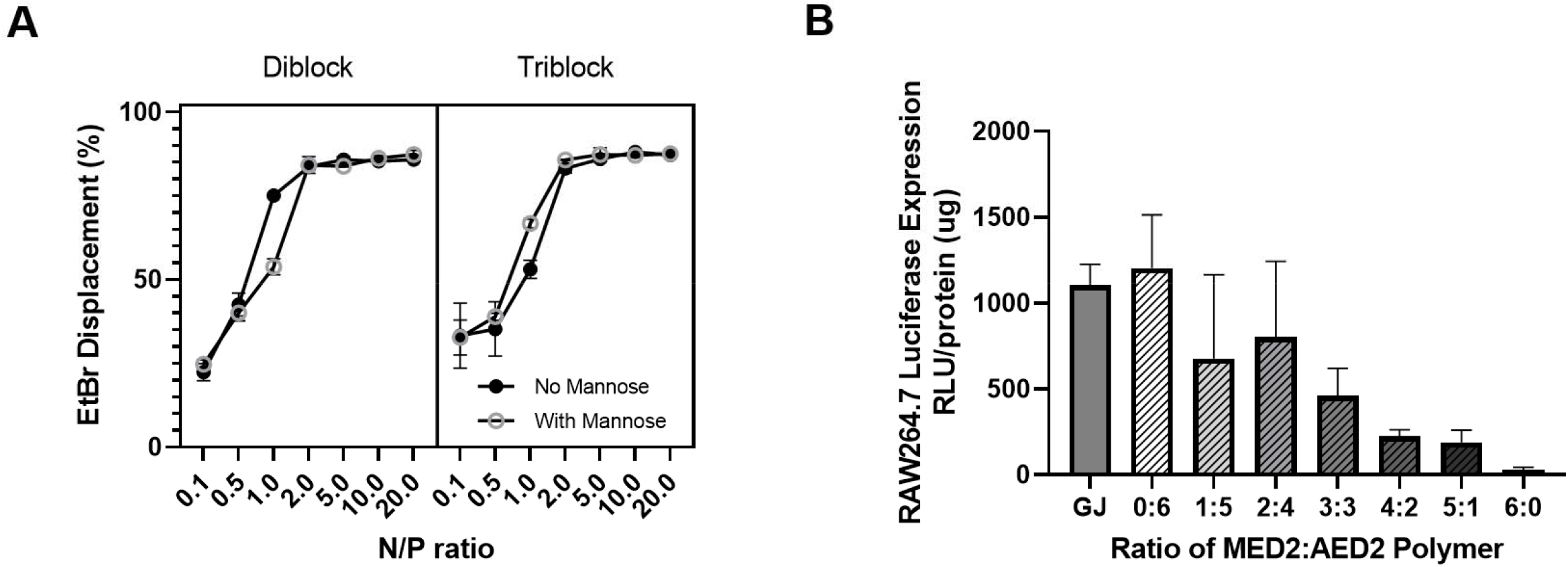
Effect of mannosylation on transfection. (A) EtBr assay of polyplexes prepared with luc-pDBA and AED2, MED2, AED3, and MED3 at various N/P ratios from 0.1 to 20. EtBr displacement (%) represent amount of EtBr displaced by polymer, or a representation of the binding between the polymer and pDNA. All polyplexes displace majority of EtBr at N/P 2 or higher. (B) Decreasing ratio of mannosylated polymer MED2 compared to ED2 shows antagonistic effect on transfection of RAW264.7 macrophages. Ratio of polymers mixed prior to forming polyplexes at N/P ratio 10.

### 2.3. Non-targeted diblock copolymer-based polyplexes transfect macrophages comparably to GeneJuice

All polyplexes were screened *in vitro* for transfection efficiency. Transfection assays allowed us to compare the cationic blocks between polymers **DMD2** (pMeOx_70_-pMestOx(DET)_50_-DBCO) or **DMT2** (pMeOx_70_-pMestOx(TREN)_50_-DBCO) which have either DET or TREN cationic moieties. In these experiments the copolymers were complexed with luc-pDNA and the resulting polyplexes were applied to IC21 macrophages for 24 hours prior to analyzing bioluminescence of the cell lysates. The **DMD2** based polyplexes transfected cells significantly better than **DMT2** based polyplexes at both N/P 10 (p<0.01) and N/P 20 (p<0.0001) **(Supplementary Figure S8 A)**. Therefore, we selected DET over TREN as a cationic moiety in the polycation in further experiments. Copolymers differing in hydrophilic MeOx or EtOx block, **DMD2** (pMeOx_70_-pMestOx(DET)_50_-DBCO) and **DED2** (pEtOx_70_-pMestOx(DET)_50_-DBCO), were also compared in *in vitro* transfection. **DED2** was significantly better at transfecting IC21 cells than **DMD2** at both N/P 10 (p<0.0001) and N/P 20 (p<0.0001**) (Supplementary Figure S8 B)**. These head-to-head comparisons led us to choose the cationic block DET and the hydrophilic block EtOx for subsequent polymer design.

The transfection efficiency of polyplexes made from **AED2, MED2, AED3**, and **MED3** were subsequently tested in RAW264.7 macrophages and bone marrow derived macrophages (BMDM). These copolymer-based polyplexes were non-toxic to cells even at higher concentrations used (high N/P ratios for polyplexes) **(Figure 3D)**. The **AED2**-based polyplexes outperformed the polyplexes made using other polymers in both RAW264.7 and BMDM transfection at both N/P ratios 10 and 20 **(Figure 4A)**. Surprisingly, the mannosylated **MED2**-based polyplexes performed significantly worse than its non-mannosylated **AED2**-based counterparts at both N/P 10 (p<0.01) and N/P 20 (p<0.0001) (Figure 4A). Polyplexes made with **AED2** at N/P 10 and 20 transfected RAW264.7 macrophages similarly to the commercial transfection reagent GeneJuice (n.s) (Figure 4A). A similar trend was seen in BMDMs; **AED2**-based polyplexes at N/P 20 performed comparably to the positive control, GeneJuice, although the overall levels of luciferase reporter gene expression normalized to the total protein were much less than those in RAW264.7 macrophages **(Figure 4B)**. Neither of the mannose-free or mannosylated triblock copolymers, **AED3** and **MED3**, transfected either cell type despite forming complexes with small size and narrow PDI at high N/P ratios. Interestingly there appeared to be an inverse correlation between the transfection efficacy of the polyplexes and the trend in their zeta potential **(Supplementary Figure S7)**. This trend is most noticeable for RAW264.7 macrophages at N/P 20.

### 2.4. Mannosylation of diblock and triblock polymers inhibits internalization and transfection

To better understand the effect of mannose on the transfection, the ratio of mannosylated to non-mannosylated diblock copolymer was varied and tested in RAW264.7 macrophage transfection. RAW264.7 cells were chosen as they showed presence of the mannose receptor CD206 which we additionally confirmed by flow cytometry **(Supplementary Figure S9)**.^[21]^ As the ratio of **MED2:AED2** increased, the transfection in RAW264.7 cells steadily decreased **(Figure 5B)**. Polyplex uptake and effect of mannose on uptake was further tested using confocal microscopy. Confocal imaging revealed that all polyplex formulations, except **MED3**-based polyplex, at N/P ratio 20 **(Figure 6)** were internalized in the cells as shown by clear Cy5 signal in Cy5 channel and merged channel. Polyplexes appear to be localized in or near lysosomes rather than dispersed in the cytoplasm. Cy5 signal was not seen inside the nucleus of any treatment group at this 24-hour timepoint. The confocal imaging data were quantified as an average Cy5 signal intensity per cell nucleus **(Supplementary Figure S10)**. To further quantify uptake, Cy5 signal was measured with flow cytometry which revealed similar results **(Supplementary Figure S11)**. Notably, the results of both quantifications appeared to correlate with the transfection results. The greatest uptake of the luc-pDNA was observed for GeneJuice transfection system and AED2-based polyplexes that also displayed the best transfection results. The uptake of the luc-pDNA in the AED3**-**based polyplexes was nearly three times less based on the confocal image quantification. Attachment of mannose residues to both diblock and triblock copolymers decreased the uptake of luc-pDNA in each case and was negligible with the MED3-pDNA polyplex, which was also inactive based on the gene expression study.

**Figure 6.**
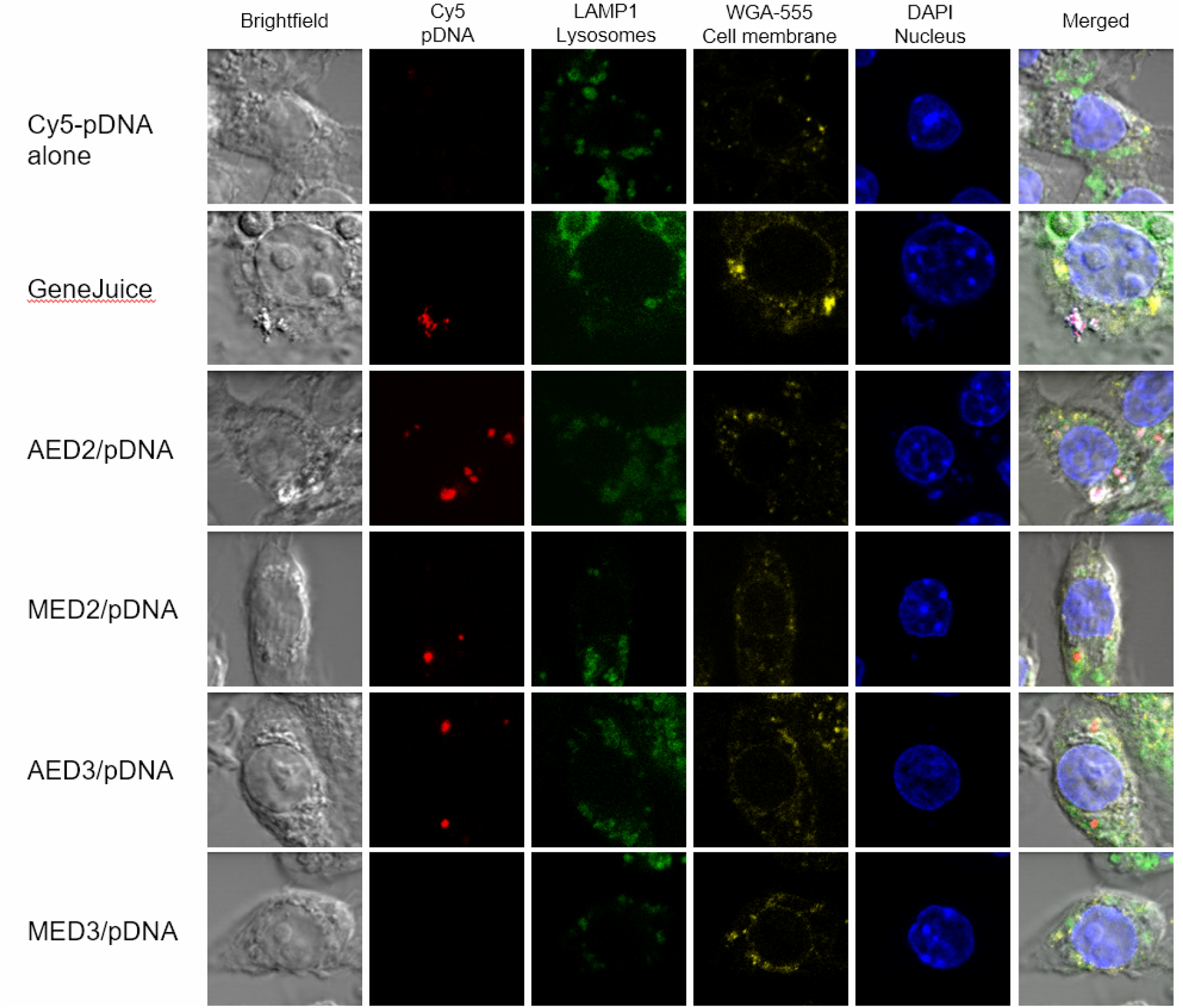
Confocal imaging of polyplex uptake in RAW264.7 macrophages. Images taken after 24-hour treatment with polyplexes. Polyplexes were prepared at N/P 20 with controls being Cy5-pDNA alone, and GeneJuice. Cellular compartments such as the cell membrane (WGA-555), lysosomes (LAMP1), and nucleus (DAPI) were stained.

## 3. Discussion

We have continued our efforts in the investigation of poly(2-oxazolines) as a viable alternative to PEG-based transfection polymers.^[22]^ Due to its biocompatibility and ability to conjugate reactive side groups, POx is a promising candidate for replacing PEG in the gene delivery applications.^[12,23–25]^ POx has been recently utilized in an assortment of biomedical applications from hydrogels to solubilizing hydrophobic drugs at high capacities.^[20,24]^ POx is also less sensitive to oxidative degradation compared to PEG.^[26]^ Out of the many POx monomers, we employed hydrophilic MeOx and EtOx, with pMeOx being slightly more hydrophilic than pEtOx. Previous findings showed that MeOx-based polymers had lower serum protein binding compared to PEG-based polyplexes, which could result in a longer circulation time *in vivo*.^[26]^ Despite the many advantages of POx which lend themselves to being advantageous for transfection, few have developed or characterized such systems for pDNA delivery. In the present work, we sought to optimize a targeted cationic POx block copolymer for efficient transfection of macrophages which are crucial immune cells in the progression of many cancers such as breast cancer. We tested the design of four polymer block components: non-ionic hydrophilic block, cationic block, hydrophobic block, and targeting moiety. We designed cationic groups to enhance transfection efficacy and developed a targeting moiety synthesis strategy to enhance uptake via the MMR. The cationic block length was kept at 50 to maintain consistency during comparison of the various polymers. We also explored the introduction of a hydrophobic block to further improve stability and complexation with luc-pDNA for future *in vivo* experiments. This study finds that a novel PEG-free POx-based pDNA delivery system is effective at transfecting a variety of macrophages including immortalized cell lines and even primary cells.

Important for designing an optimized system for gene delivery are 1) the chemical composition of the nonionic block, 2) the structure of the cationic block side chains, and 3) the hydrophilicity of the block copolymer, all of which can affect the interaction between polyplexes and cell membranes.^[27,28]^ When comparing the transfection efficiency of polyplexes made with polymers containing either MeOx or EtOx hydrophilic block, we found that the EtOx-based polyplexes outperformed those made with most hydrophilic MeOx. Probably, the MeOx shell of the corresponding polyplex was too hydrophilic that it masked not only the binding of the serum proteins as we have previously shown, but also hindered the polyplex interaction with cells.^[22]^ The effect of the nanoparticle hydrophilic shell structure on their uptake in macrophages has been shown for liposomes coated with PEG and hyperbranched polyglycerol.^[29]^ Our previous study found that pMeOx and pEtOx-conjugated protein internalized at higher rates compared to PEGylated protein in CATH.a neuronal cells.^[30]^ This study observed that EtOx-based conjugates are internalized at a ∼4 to ∼7 times faster rate compared to MeOx-based conjugates, which supports why EtOx-based polyplexes transfect cells more efficiently than MeOx-based polyplexes.^[30]^ Since both MeOx and EtOx-based polyplexes formed complexes of similar size, it not likely to be responsible for the difference in IC21 transfection. Overall, the EtOx monomer was chosen for subsequent studies as it showed greatest transfection efficacy.

In the following studies, we focused on comparing the cationic blocks with different side chains, DET and TREN, to determine which one is more efficient in transfecting macrophages. Previously, DET was used in PEG-containing transfecting polymers.^[31,32]^ Kataoka first reported the cationic block copolymers with DET-modified poly(L-aspartic acid) block as good transfecting agents that exhibited a proton sponge effect facilitating the pDNA delivery to cells and were safe to cells *in vitro*.^[33,34]^ TREN is another cationic moiety which is commonly used in lipid-based transfection systems due to its branched structure which allows for efficient condensation of genetic material.^[35–37]^ Both cationic moieties, DET and TREN, were chosen based on good biocompatibility, and different charge densities of linear versus branched structures which could impact the complexation with pDNA.^[34]^ As expected, TREN-based block copolymers formed tighter complexes perhaps due to the difference in the charge density compared to DET-based block copolymers.^[35,38]^ Despite forming a more stable complex, TREN-based polyplexes transfected macrophages poorly compared to DET-based polyplexes. As previously reported, tightly bound polyplexes are not able to release their genetic cargo and therefore are worse transfection agents.^[27,28,39]^ Notably, the DET-containing copolymers exhibited buffering capacity between pH 5.7 to 7.0, while the TREN-containing copolymers did not. Since the most widely accepted theory of endosomal escape of nucleic acids relies on the ability to attract protons as stated in the proton sponge theory, the DET side chain is a good candidate for nucleic acid delivery into the cell.^[40–43]^ With both diblock and triblock copolymers, DET proved to be a cationic moiety capable of forming well-defined polyplexes with luc-pDNA leading us to choose it as the optimal cationic block.

Two synthetic click chemistry strategies were used to introduce the targeting moiety to develop the least toxic clickable system for future *in vivo* success. For the cell transfection studies mannose was conjugated via CuAAC rather than copper-free AAC due to a greater mannose conjugation with the CuAAC method. Though the CuAAC method uses copper as a catalyzing reagent, the mannosylated polymers did not show toxicity. Mannosylated copolymers based on DET and EtOx were expected to increase transfection by increasing targeting to macrophages and therefore also increasing uptake. However, both transfection and uptake in macrophages were hindered when using mannosylated polyplexes made from diblock and triblock polymers. Despite a lack of toxicity, and mannose conjugation extent at 37 and 31% for MED2 and MED3 respectively, mannosylation did not improve transfection. Even when varying the ratio of MED2 to non-mannosylated AED2 in polyplex formation, the greater amount of AED2 resulted in increased transfection of RAW264.7 macrophages. Blakney et al. reported that when PEI was modified with mannose, the transfection with small activating RNA (saRNA) in HEK293 cells was decreased, potentially due to steric hindrance of mannose.^[18]^ This group also reported that as amount of mannose moieties attached to PEI was increased, the transfection decreased, which is a similar trend found in the present study.^[18]^ When also comparing EtBr displacement, mannosylated polymers did not displace differently compared to non-mannosylated counterparts meaning that they did not differ much in their binding to pDNA. The localization quantification shows that the cellular uptake of cy5-luc-pDNA is decreased in polyplexes made with mannosylated polymers at N/P ratio 20. Thus, non-mannosylated diblock and triblock polyplexes had greater uptake compared to their mannosylated counterparts. When analyzing the internalization of Cy5-pDNA by flow cytometry, the uptake trend was similar to confocal imaging quantification suggesting that mannosylated polyplex transfection is being hindered during uptake.

Currently, polyplexes of various sizes are believed to enter the cell through various endocytosis pathways.^[44,45]^ Notably the mannosylated copolymer-based polyplexes had similar size by DLS but their uptake in macrophages was inhibited compared to non-targeted polymers. Endocytosis is also governed by shape of particles. The slightly elongated shapes of polyplexes made with mannosylated copolymers could contribute to decreased uptake as Skirtach et al. reports that high-aspect ratio particles result in slower and overall decreased uptake compared to spherical particles due to the forces generated at the interaction between cell and particle.^[41,44]^ Therefore, the elongated worm-like shape of mannosylated polyplexes can result in decreased uptake, though there is no clear consensus in the literature. Non-mannosylated triblock AED3-based polyplexes also had both low uptake and poor transfection. Since triblock polyplexes had a hydrophobic core, this could cause the formation of complexes which are too stable for releasing pDNA cargo. Therefore, uptake is an indicator of transfection success, and mannosylation on these diblock and triblock copolymers interferes with that process. As mannosylation has been previously reported as an enhancer of internalization, it is surprising that mannose conjugation did not improve uptake or transfection in the present study. Though more studies need to be done to confirm the true cause of this inhibition, potential aspects to study include flexibility of polymer chains, surface charge at various points during endocytosis, timing of uptake, and incomplete click chemistry.

## 4. Experimental sections

### 4.1. Materials

Monomers were purchased from Sigma-Aldrich (St. Louis, MO). Polymers were synthesized in acetonitrile (ACN) with the initiators propargyl *p*-toluenesulfonate or p-toluenesulfonic acid methyl ester using the following purified monomers: 2-ethyl-2-oxazoline (EtOx), 2-methyl-2-oxazoline (MeOx), 2-methoxy-carboxyethyl-2-oxazoline (MestOx), and 2-isopropyl-2-oxazoline (iPrOx). Cationic modifications were made with diethylenetriamine (DET) or tris(2-aminoethyl)amine (TREN). Polymers were terminated with either 3-Amino-1-[(5-aza-3,4:7,8-dibenzocyclooct-1-yne)-5-yl]-1-propanone (dibenzocyclooctyne-amine or DBCO-amine) or piperidine. Alpha-Mann-TEG-N_3_ (Iris Biotech, Marktredwitz, Germany) (mannose) was conjugated as a targeting moiety to the alkyne via click chemistry. The 2-(p-toluidino)-6-naphthalene sulfonic acid (TNS) was from Millipore (Sigma). Polyplexes were formed using polymers and gWIZ luciferase-encoding plasmid (luc-pDNA) (Gene Therapy Systems, San Diego, CA) and expanded using Plasmid Giga Kit (Qiagen, Hilden, Germany). Polyplexes were mixed with 6X orange loading dye (Thermo Fisher Scientific, Waltham, MA) prior to running agarose gel electrophoresis in 1X TAE buffer to confirm complexation. Ethidium Bromide (EtBr) (Thermo Fisher Scientific, Waltham, MA) was added to the agarose gel for visualizing the luc-pDNA under UV illumination. Cells were transfected with commercially available non-lipid transfection reagent GeneJuice as the positive control using the manufacturer’s protocol (EMB Millipore Novagen, Madison, WI). Transfection efficacy *in vitro* was analyzed with dual-assay reporter kit (Promega, Madison, WI) and normalized with Pierce™ BCA Protein Assay Kit (Thermo Fisher Scientific, Waltham, MA). The pDNA was covalently labeled with Cy5 using the Label IT™ Nucleic Acid Labeling Kit (Mirus Bio, Madison, WI). Internalization imaging was performed in Nunc™ Lab-Tek™ II Chambered Coverglass (Thermo Fisher Scientific, Waltham, MA). Uptake was quantified by flow cytometry using Zombie Violet™ Fixable Viability Kit (BioLegend, San Diego, CA). Cytotoxicity was evaluated with CCK-8 assay (Dojindo, Rockville, MD).

### 4.2. Methods

#### 4.2.1. Synthesis of block copolymers

##### Synthesis conditions

Here and below all copolymers were synthesized in ACN via sequential LCROP carried out in optimal glovebox conditions with H_2_O and O_2_ levels always maintained below 10 ppm and 20 ppm, respectively. ^1^H-NMR spectra were recorded on an INOVA 400 (Agilent Technologies, Santa Clara, CA) at room temperature. The spectra were calibrated using the solvent signals (D_2_O 4.80 ppm).

##### Synthesis of DBCO-containing block copolymers

The DBCO-containing block copolymers, DBCO-pMeOx_70_-pMestOx(DET)_50_ (DMD2), DBCO-pEtOx_70_-pMestOx(DET)_50_ (DED2), and DBCO-pEtOx_70_-pMestOx(TREN)_50_ (DMT2), were synthesized as follows. The reaction was initiated by p-toluenesulfonic acid methyl ester (0.238 mmol, 1 eq.) followed by sequential polymerization of MestOx (11.89 mmol, 50 eq.) and either MeOx (16.67 mmol, 70 eq.) or EtOx (16.67 mmol, 70 eq.). Monomers were sequentially added to the reaction mixture dropwise and stirred at 80 °C for two days for the first and second blocks. The reaction was terminated by DBCO-amine (0.714 mmol, 3 eq.). The resulting polymer precursors **DMD2** (MeOx block) or **DED2** (EtOx block) were modified with DET or TREN by stirring each mannosylated polymer (20 mg) in a DET or TREN solution (2 mL) at 40 °C for three days. Excess DET was purified by dialysis and final polymers **DMD2** and **DED2**, modified by DET, and **DMT2**, modified by TREN, were collected by lyophilization, and stored at –20 °C.

##### Synthesis of alkyne-containing block copolymers

The alkyne-containing block copolymers, pEtOx_70_-pMestOx(DET)_50_ (AED2) and Alkyne-pEtOx_50_-pMestOx(DET)_50_-piPrOx_20_ (AED3), were synthesized as follows. The reaction was initiated by propargyl *p*-toluenesulfonate (0.238 mmol, 1 eq.) in acetonitrile (5 mL) followed by sequential polymerization of EtOx (**AED2**: 16.67 mmol, 70 eq., **AED3**: 11.89 mmol, 50 eq.,) and then MestOx (11.89 mmol, 50 eq.), and, in case of triblock copolymer, iPrOx (4.76 mmol, 20 eq,). All monomers were added to the reaction media dropwise and the reaction was carried upon constant stirring at 80 °C overnight for the first block, three days for the second block, and overnight for the third block. The reactions were terminated by piperidine (0.714 mmol, 3 eq.) which was added after completion of either second or third block. After each polymerization step and reaction termination, the precursor polymers (diblock, AE2, or triblock, AE3) were characterized with ^1^H NMR. Acetonitrile was removed in vacuo. Finally, precursors **AE2** or **AE3** were either conjugated to mannose and then modified with DET, or just modified with DET. For non-mannosylated polymers, excess DET was added to precursors **AE2** and **AE3** by stirring dry polymer (20 mg) in a DET solution (2 mL) at 40 °C for 3 days. Excess DET was purified by dialysis against 3.5 kDa MWCO membrane in 0.01N HCl overnight followed by dialysis in DI water for 2 days. Diblock **AED2** and triblock **AED3** were lyophilized and stored at –20 °C.

##### Mannose conjugation to the block copolymers

The mannose-pEtOx_70_-pMestOx(DET)_50_ (**MED2**) and mannose-pEtOx_50_-pMestOx(DET)_50_-piPrOx_20_ (**MED3**) were prepared as follows. Alpha-Man-TEG-N3 (mannose) was conjugated to the ethyl oxazoline block via CuAAC click chemistry. Stock mannose was diluted with DI water to 200 mg/mL and stock solutions were kept at -20 °C. After reconstituting **AE2** or **AE3** (50 mg) in DI water, mannose (16.9 mg) was added, and the solution was stirred for 5 min. Next, CuSO_4_*5H_2_O (2.4 mg) and sodium ascorbate (3.0 mg) were added sequentially. Volume of final solutions was kept at 1 mL and stirred at room temperature (RT) overnight. Next, solutions were dialyzed against DI water in a 3.5k MWCO membrane for 2 days. **MED2** and **MED3** were then lyophilized and analyzed using NMR. Mannose ligand conjugation yield: 37% for **MED2**, 31% for **MED3**; ^1^H NMR (400 MHz, D_2_O, 25 °C).

#### 4.2.2. Polymer characterization experiments

##### Plasmids

The gWIZ™ high expression vector encoding the reporter gene luciferase (luc-pDNA) was used throughout the study. The plasmid was expanded with the Plasmid Giga Kit following the supplier’s protocol and stored at -20 °C until needed.

##### Polyplex preparation

Polyplexes at various N/P ratios were prepared using polymers and gWIZ™ luciferase-encoding pDNA. Polymers were serially diluted with 10 mM HEPES buffer according to desired N/P ratio and briefly mixed with a fixed amount of luc-pDNA using vortex mixer. Polyplexes were incubated at RT for 30 min. For further characterization at physiological conditions, 3M NaCl was added to polyplex solutions to reach final concentration of 150 mM NaCl. Those solutions were then incubated at 37 °C for 60 min. For characterization studies, aliquots of polyplex solution were taken after the first or second condition and analyzed by DLS, NTA, zeta potential, or gel electrophoresis. For transfection studies in 24-well flat bottom plates, polyplexes were prepared at RT for 30 min and mixed with serum-free media prior to application to the cells.

##### Dynamic light scattering

Z-average hydrodynamic diameter and polydispersity index (PDI) were determined by dynamic light scattering (DLS) using a Malvern Zetasizer (Malvern Instruments, Westborough, MA). Samples for DLS were prepared in 10 mM HEPES buffer (50 uL) and measured in triplicate with a minimum of 10 runs per measurement per sample. Measurements were taken either after 30 min incubation at RT or after 30 min incubation at RT followed by 60 min incubation at 37 °C.

##### Zeta potential measurement

Polyplexes were prepared at N/P 20 and incubated at RT for 30 min. Samples were diluted with DI water (total volume 1 mL) and measured on a Malvern Zetasizer (Malvern Instruments, Westborough, MA).

##### Luc-pDNA incorporation (agarose gel electrophoresis)

To confirm luc-pDNA complexation with polymers, gel electrophoresis was performed. Gel loading dye (orange, 6X) was used for each sample. Experimental samples were compared to naked luc-pDNA alone. Samples were loaded onto 1% agarose gel in 1X TAE buffer and run at 100V for 45 min. Gels were imaged by UV illumination.

##### EtBr displacement assay

To determine the relative binding affinity between luc-pDNA and polymers, ethidium bromide (EtBr) displacement was measured. Briefly, EtBr was diluted to 2 μg/mL and mixed with luc-pDNA to a final luc-pDNA concentration of 33 μg/mL. Polyplexes were formed in an opaque 96-well black plate at various N/P ratios ranging from 0.1 to 20. EtBr displacement was quantified by measuring fluorescence at Ex/Em 520/590 nm emission using a SpectraMax M5 plate reader (Molecular Devices, San Jose, CA). Relative fluorescence (%) was calculated by subtracting EtBr alone background fluorescence from each experimental sample and normalizing to fluorescence of a control solution containing only luc-pDNA and EtBr.

##### Transmission electron microscopy (TEM)

All TEM images were obtained on a Talos F200X S/TEM microscope (Thermo Fisher Scientific, Waltham, MA). Polyplex samples prepared at N/P 20 were applied to 300 mesh carbon-coated copper grids and stained with 4% uranyl acetate prior to imaging (Ted Pella, Redding, CA). Excess sample was blotted gently and allowed to air dry prior to imaging.

##### TNS assay

A TNS assay assessed the surface charge and apparent pKa of polymers with DET or TREN side chains by measuring the fluorescence intensity change in solutions of polyplexes mixed with fluorescent TNS over a range of pH from 4.0 to 7.4. Polyplexes were prepared in 10 mM citrate buffer (300 μL), containing 150 mM NaCl, at pH 4.0, 5.0, 6.0, and 7.4 and mixed with 3 μL of 6 mM TNS using vortex mixing. Fluorescence intensity of samples was measured in triplicate with 100 μL volume per well in a 96-well black plate at Ex/Em 325/435 nm using a SpectraMax M5 plate reader (Molecular Devices, San Jose, CA). Fluorescence intensity was normalized to fluorescence at pH 7.4.

##### Acid–base titration assay

The cationic block copolymers were dissolved in 10 mM HCl-containing saline at the cationic repeating unit base-molar concentration of 3 mM (the base-molar concentration represents the polymer molar concentration multiplied by the degree of polymerization of the cationic block). Initial pH 2 was recorded and small amounts of 0.1 M NaOH were added while measuring pH after each addition until reaching pH 12. To analyze the buffering capacity, the change in dOH^-^ was divided by dpH for each measurement in the titration. The resulting value indicates how much OH^-^ is needed to increase pH.

#### 4.2.3. Cellular experiments

##### Cell culture

RAW264.7 macrophages were cultured in DMEM media supplemented with 10% FBS and 1% penicillin/streptomycin (p/s). IC21 macrophages were cultured in RPMI media supplemented with 10% FBS and 1% p/s. Bone marrow-derived macrophages (BMDM) (129/sv background) were isolated from the femur of a mouse. The monocytes were cultured for 10 days in DMEM media supplemented with 10% FBS and MCSF-containing media obtained from L929 cells. BMDMs were used on Day 10. All cell cultures were maintained at 37 °C and 5% CO_2_.

##### Cytotoxicity

Polyplexes were prepared at N/P ratios 10 and 20 as previously described. RAW264.7 cells were seeded at 10^4^ cells/well in a 96-well plate. Prior to transfection, serum-containing media was replaced with serum-free DMEM. Cells were then treated with media, luc-pDNA alone, GeneJuice, or polyplexes for 24 hours with each well containing 0.25 μg luc-pDNA. After incubation, fresh serum-containing DMEM was applied containing 10% CCK-8 solution. Absorbance was read at 450nm after 1 hr.

##### in vitro transfection

Polyplex formulations were prepared at N/P ratios 10 and 20. IC21, RAW264.7, or primary BMDM were seeded in a 24-well plate. After reaching 70% confluency, cells were treated with luc-pDNA alone, GeneJuice (positive control), or polyplexes for 24 hours with each well receiving luc-pDNA (1 μg). After treatment, cells were rinsed once in DPBS (500 μL) and lysed in 1X cell culture lysis buffer (100 μL) for 45 min on a shaker plate at RT. Lysates were collected and either immediately analyzed for luciferase activity or stored at -80 °C for further analysis. Final luciferase activity was either reported as RLU or further normalized by total protein in cell sample (RLU/ μg total protein).

##### Bioluminescence analysis of transfected cell lysates

After transfection, cell lysates were analyzed for luciferase activity using a luciferase reporter assay following the manufacturer protocol. To normalize luciferase activity results per well, total protein was quantified using the Pierce™ BCA assay kit in a 96-well plate. Final luciferase activity was calculated as luciferase/total protein (RLU/μg total protein). All samples were measured in triplicate.

##### Mannose receptor presence verification via flow cytometry

Macrophages RAW264.7, IC21, or BMDMs were analyzed for mannose receptor (MMR; CD206) presence. Samples were analyzed on an LSR II or LSR Fortessa cytometer (BD Biosciences, San Jose, CA). A minimum of 10,000 events were recorded. Results are shown as a histogram of CD206+ intensity relative to the unstained sample intensity.

##### Uptake via confocal microscopy

In an 8-well chambered coverglass slide, RAW264.7 macrophages were treated for 24 hours with polyplexes formed at N/P ratio 20 with Cy5-labeled luc-pDNA. Each well was treated with a total of 1 μg Cy5-pDNA. Polyplex uptake was compared to controls such as cells alone, luc-pDNA alone, and GeneJuice transfection reagent. The following cellular compartments were stained: lysosomes (LAMP1), cell membrane (WGA-555), and nuclei (DAPI). Images were taken at 40X magnification on a Zeiss LSM710 (Carl Zeiss AG, Oberkochen, Germany) inverted laser scanning confocal microscope.

##### Uptake via flow cytometry

In a 6-well plate, RAW264.7 macrophages were treated for 24 hours with polyplexes formed at N/P ratio 20 with Cy5-labeled luc-pDNA. Each well was treated with a total of 2 μg Cy5-pDNA. Samples were stained with Zombie Violet Live/Dead dye and the 405 nm and 633 nm laser were used to excite fluorophores. Fluorescence data was collected on an LSR Fortessa cytometer (BD Biosciences, San Jose, CA). One sample represents one well. A minimum of 10,000 events were recorded. Results are shown as a histogram of Cy5+ intensity relative to the unstained sample intensity.

##### Statistical analysis

Experimental samples (n=3∼4) were compared using either student’s t-test or one-way ANOVA with multiple comparisons. Statistical significance was obtained using GraphPad Prism 9 with *p<0.05, **p<0.01, ***p<0.001, ****p<0.0001.

## 5. Conclusion

This study designed and characterized a POx-based platform for transfecting macrophages with pDNA. Optimal diblock and triblock configurations for highest transfection efficiency consisted of a hydrophilic EtOx block and a cationic DET moiety. The hydrophobic iPrOx block was introduced for the triblock structure which did not improve transfection. The polyplexes were uniformly sized and safe to macrophages *in vitro*. Mannosylation of polymers did not enhance the uptake or transfection of macrophages in this specific polymer design. Uptake was also affected by surface charge of complexes where the less positively charged polyplexes transfected the cells more efficiently. Polyplexes made with luc-pDNA and a diblock POx polymer consisting of a hydrophilic EtOx block and a cationic DET moiety transfected both immortalized and primary macrophages with the same efficiency as the commercial transfection reagent, GeneJuice. This study developed an efficient non-toxic PEG-free polymer, AED2, capable of transfecting macrophages with pDNA efficiently.

## Supporting Information

Supporting Information is available from the Wiley Online Library or from the author.

## Acknowledgements

This work was performed in parts at the Chapel Hill Analytical and Nanofabrication Laboratory, CHANL, a member of the North Carolina Research Triangle Nanotechnology Network, RTNN, which is supported by the National Science Foundation, Grant ECCS-1542015, as part of the National Nanotechnology Coordinated Infrastructure; the Microscopy Services Laboratory, Department of Pathology and Laboratory Medicine, supported in part by P30 CA016086 Cancer Center Core Support Grant to the UNC Lineberger Comprehensive Cancer Center; and the Nanomedicine Characterization Core facility at the UNC Center for Nanotechnology in Drug Delivery, supported by the Carolina Partnership between the UNC Eshelman School of Pharmacy and the University Cancer Research Fund. Olesia Gololobova of the UNC Nanomedicine Characterization Core facility is acknowledged for assistance in some experiments.

Dina N. Yamaleyeva performed polyplex formation, characterization, cytotoxicity, uptake and transfection experiments, prepared figures, and drafted manuscript. Naoki Makita proposed specific copolymer structures and synthetic schemes, performed copolymer characterization, polyplex formation and characterization experiments. Duhyeong Hwang proposed specific copolymer structures and synthetic schemes, performed copolymer synthesis, characterization, and chemical modification experiments. Matthew Haney performed selected cell transfection experiments. Alexander V. Kabanov developed concept and approach, supervised research, drafted manuscript. Rainer Jordan developed concept and approach.

