## Supplemental Figures for "Poly(2-oxazoline)-based polyplexes as a PEG-free plasmid DNA delivery platform"

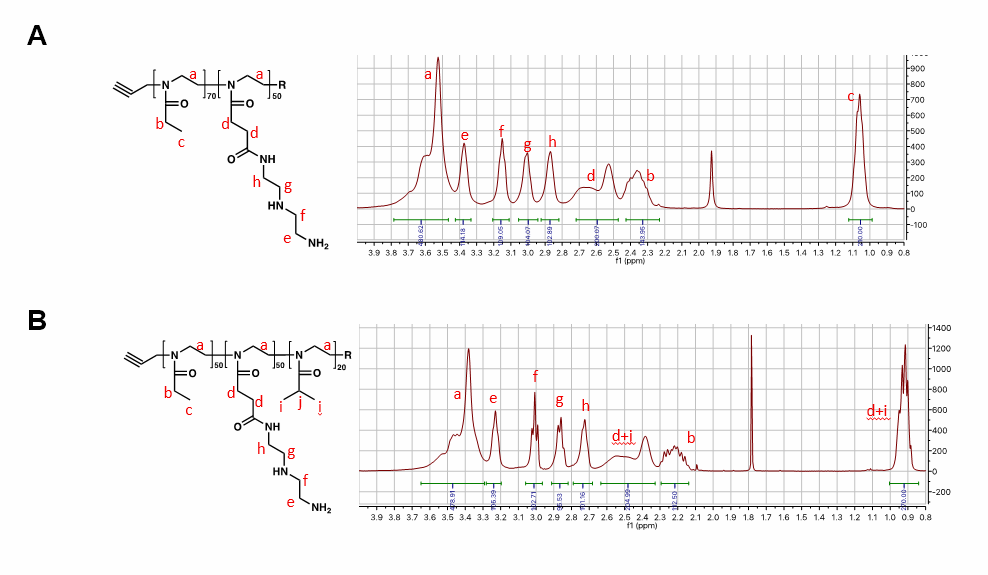

**Supplementary Figure S1.** NMR of diblock (AED2) and triblock (AED3) polymers. (A) AED2. (B) AED3.

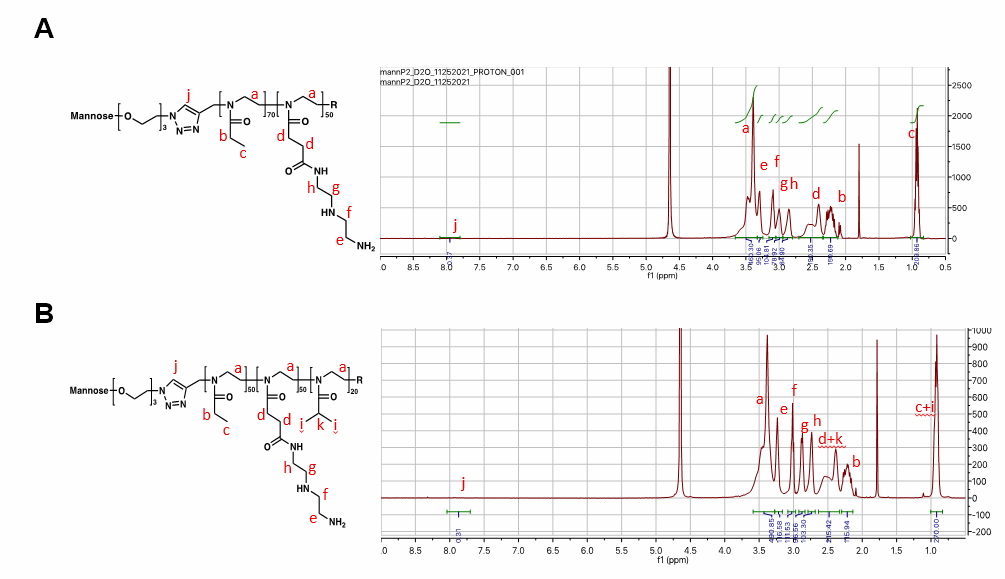

**Supplementary Figure S2.** NMR of mannosylated diblock (MED2) and triblock polymers (MED3). (A) MED2 has 37% mannose conjugation. (B) MED3 has 31% mannose conjugation.

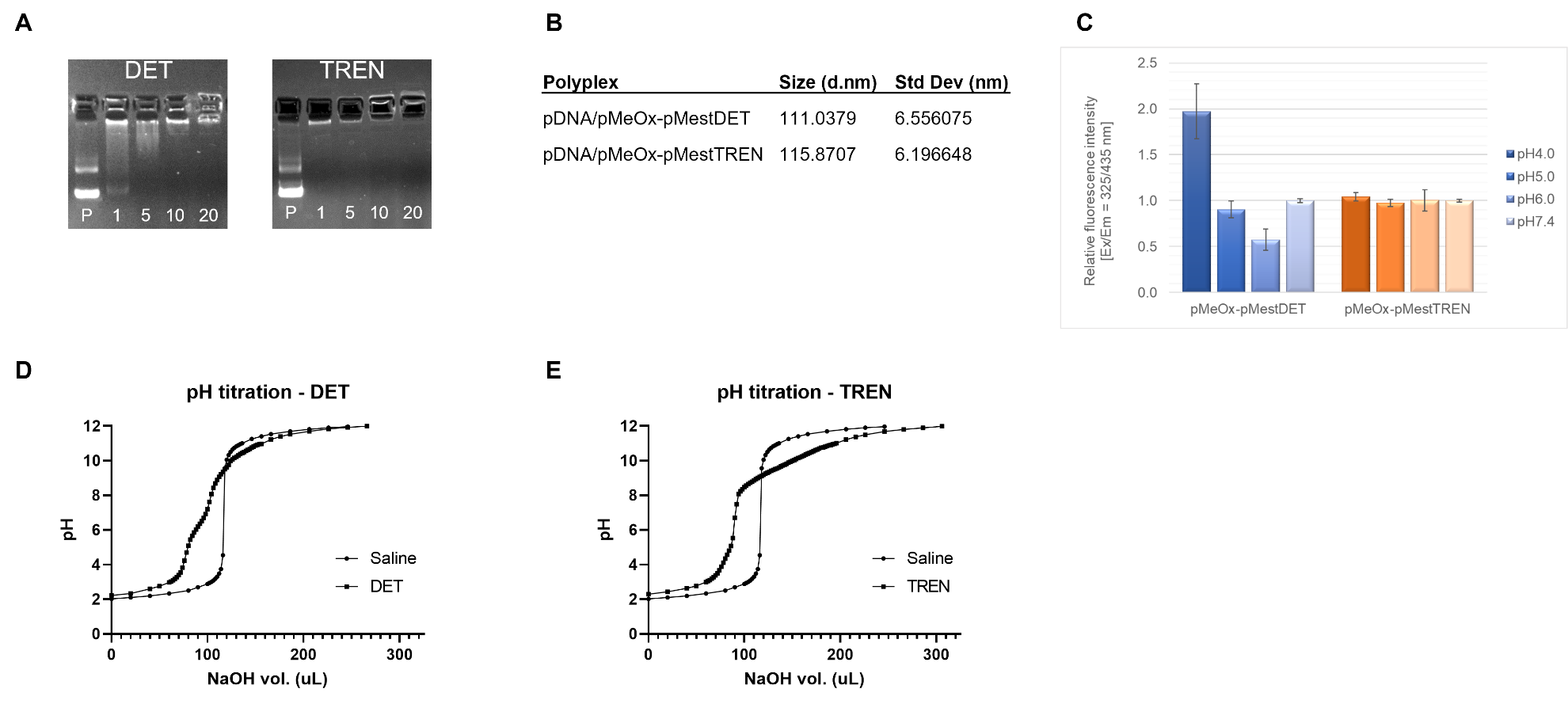

**Supplementary Figure S3.** Physicochemical characterization of cationic moieties, DET and TREN. (A) An image of agarose gel electrophoresis for pDNA complexed with DMD2 (pMeOx-pMestDET, left) and DMT2 (pMeOx-MestTREN, right) at various N/P ratios. pDNA was visualized by ethidium bromide staining. P: pDNA alone. (B) Polyplex size distribution of DMD2-pDNA and DMT2-pDNA complexes at N/P = 10 as analyzed by DLS. Effective diameter given in nm ±SD. (C) TNS assay results of pMeOx-pMestDET and pMeOx-pMestTREN. Error bars represent ± SD. (D) pH titration curves of pMeOx-pMestDET and (E) pMeOx-pMestTREN solutions.

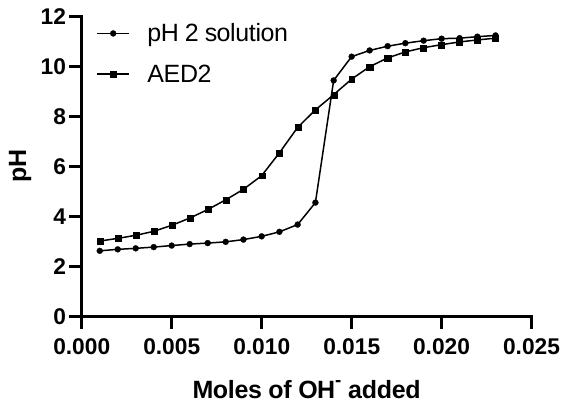

**Supplementary Figure S4**. Buffering capacity analysis of EtOx-based DET-containing block copolymer, pEtOx-pMestDET. Resulting pKa of pEtOx-pMestDET is at approximately pH 4.3 and 8.8.

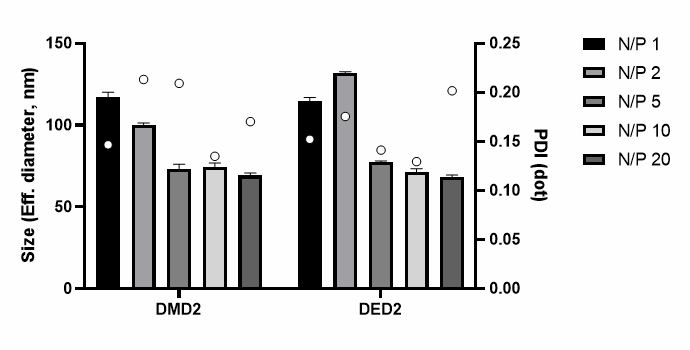

**Supplementary Figure S5.** Size comparison of polyplexes made with various hydrophilic blocks. Copolymers tested were DMD2 (pMeOx-pMestDET) or DED2 (pEtOx-pMestDET), at various N/P ratios. Size of polyplexes formed with pDNA at N/P ratios 1, 2, 5, 10, 20. Error bars represent +/- SD.

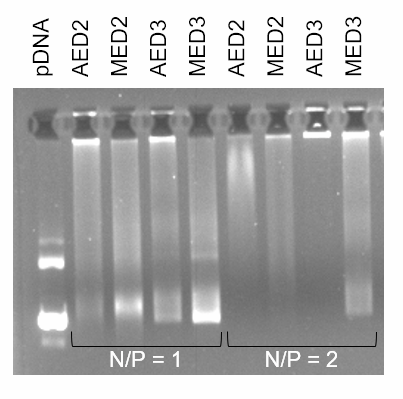

**Supplementary Figure S6.** Gel electrophoresis. Gel electrophoresis showing complexation between polymers AED2, MED2, AED3, MED3 and luc-pDNA at NP ratios 1 and 2 after 30 min incubation at RT. Polyplexes were measured on a 1% agarose gel at 100V for 45 min.

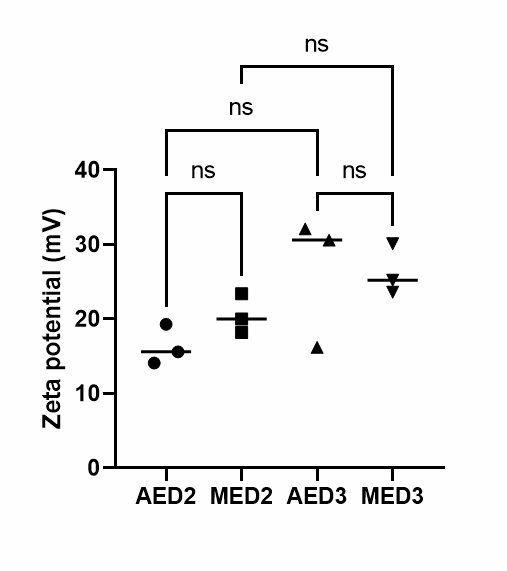

**Supplementary Figure S7.** Zeta potential of polyplexes. Polyplexes made with polymers AED2, MED2, AED3, or MED3 and luc-pDNA were prepared at N/P 20 and incubated at RT for 30 min.

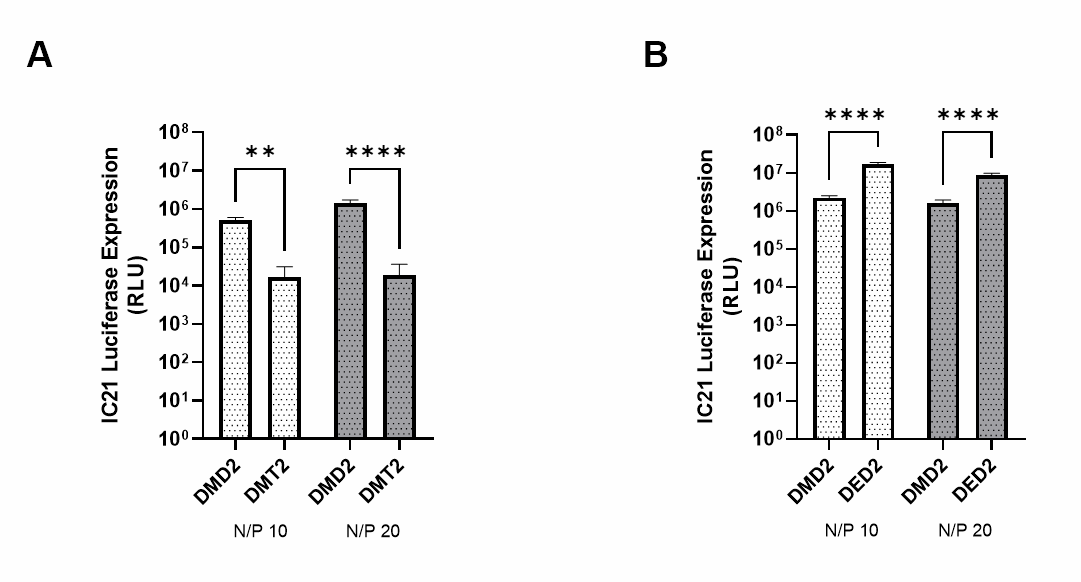

**Supplementary Figure S8.** POx-pDNA polyplex optimization via *in vitro* transfection of IC21 macrophages. (A) Cells transfected with either DMD2-based (DET) or DMT2-based (TREN) polyplexes at N/P 10 and 20. (B) Cells transfected with either DMD2-based (MeOx) or DED2-based (EtOx) polyplexes at N/P 10 and 20. *p<0.05, **p<0.01, ***p<0.001, ****p<0.0001.

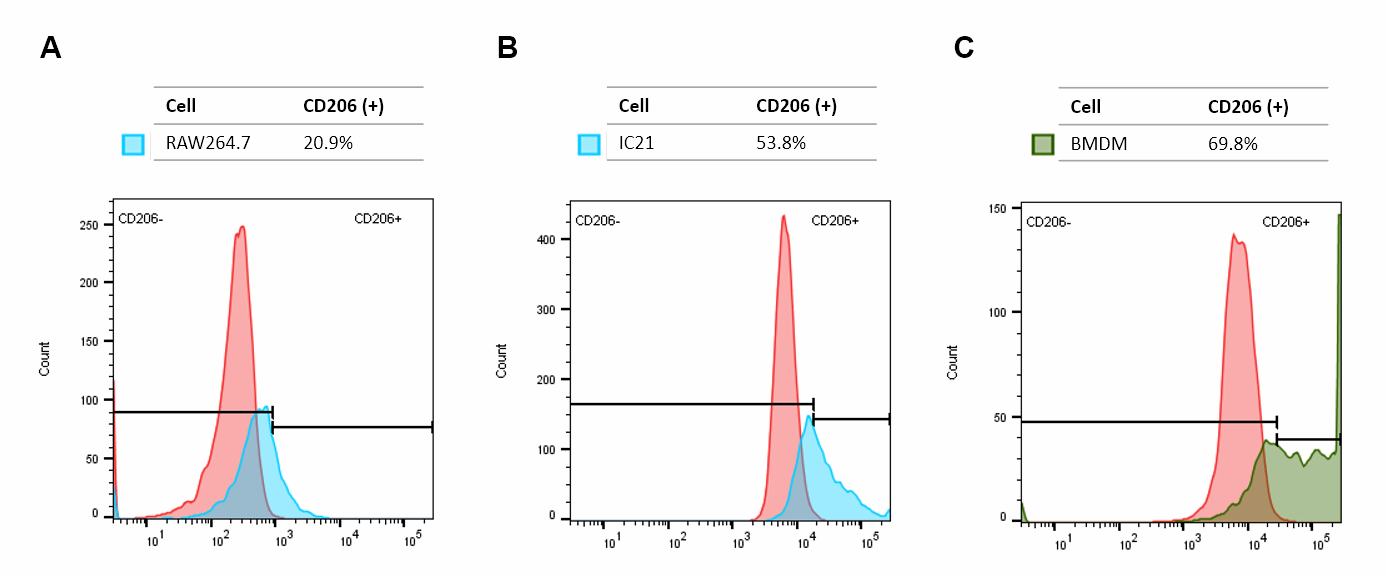

**Supplementary Figure S9**. Mannose receptor (CD206) presence analyzed via flow cytometry. CD206 presence analyzed on unstimulated cells: (A) RAW264.7, (B) IC21, and (C) BMDM. Samples compared to unstained cells (red histogram).

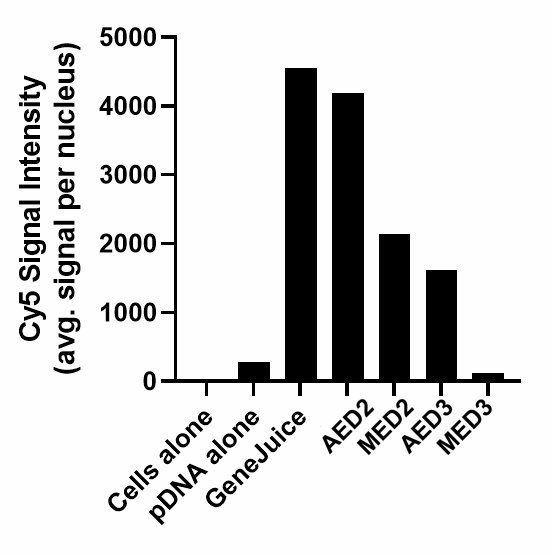

**Supplementary Figure S10.** Confocal imaging quantification of uptake. RAW264.7 macrophages were treated with polyplexes prepared at N/P 20. Cy5+ signal represents internalized Cy5-pDNA as average signal intensity per nucleus.

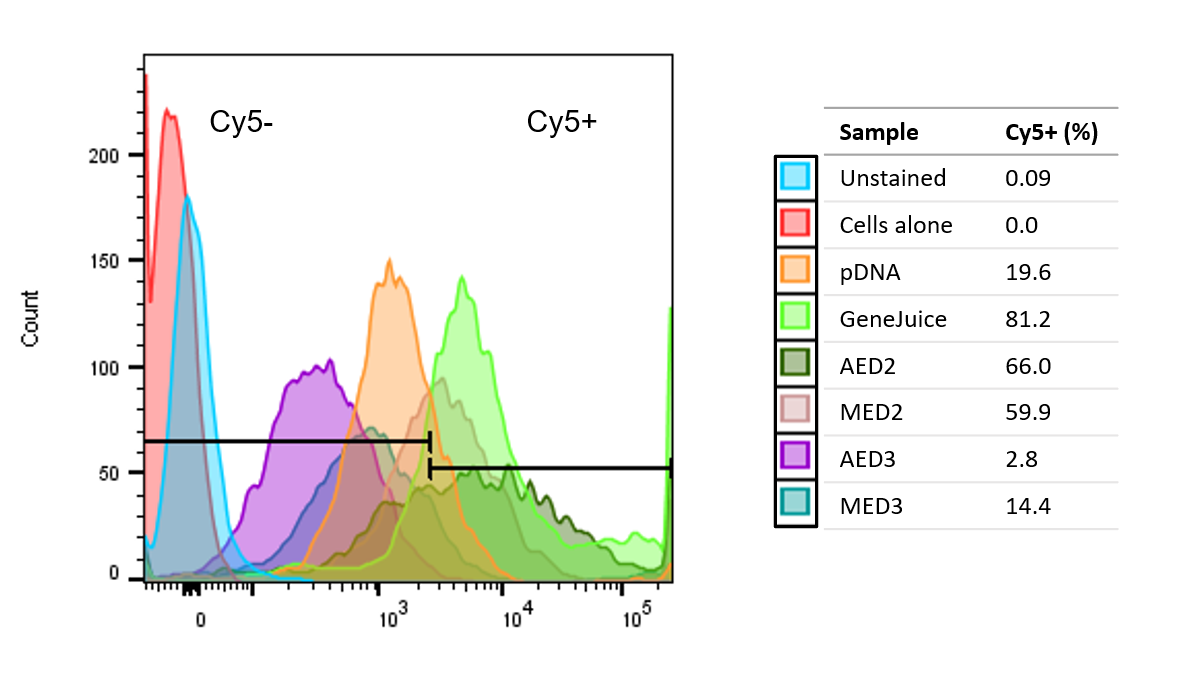

**Supplementary Figure S11.** Flow cytometry quantification of uptake. RAW264.7 macrophages were treated with polyplexes prepared at N/P 20. Cy5+ signal represents internalized Cy5-pDNA.
